# Cortical Processing of Arithmetic and Simple Sentences in an Auditory Attention Task

**DOI:** 10.1101/2021.01.31.429030

**Authors:** Joshua P. Kulasingham, Neha H. Joshi, Mohsen Rezaeizadeh, Jonathan Z. Simon

**Affiliations:** Department of Electrical and Computer Engineering, University of Maryland, College Park, Maryland; Institute for Systems Research, University of Maryland, College Park, Maryland; Department of Biology, University of Maryland, College Park, Maryland

## Abstract

Cortical processing of arithmetic and of language rely on both shared and task-specific neural mechanisms, which should also be dissociable from the particular sensory modality used to probe them. Here, spoken arithmetical and non-mathematical statements were employed to investigate neural processing of arithmetic, compared to general language processing, in an attention-modulated cocktail party paradigm. Magnetoencephalography (MEG) data was recorded from 22 subjects listening to audio mixtures of spoken sentences and arithmetic equations while selectively attending to one of the two speech streams. Short sentences and simple equations were presented diotically at fixed and distinct word/symbol and sentence/equation rates. Critically, this allowed neural responses to acoustics, words, and symbols to be dissociated from responses to sentences and equations. Indeed, the simultaneous neural processing of the acoustics of words and symbols was observed in auditory cortex for both streams. Neural responses to sentences and equations, however, was predominantly to the attended stream, and originated primarily from left temporal, and parietal areas, respectively. Additionally, these neural responses were correlated with behavioral performance in a deviant detection task. Source-localized Temporal Response Functions revealed distinct cortical dynamics of responses to sentences in left temporal areas and equations in bilateral temporal, parietal and motor areas. Finally, the target of attention could be decoded from MEG responses, especially in left superior parietal areas. In short, the neural responses to arithmetic and language are especially well segregated during the cocktail party paradigm, and the correlation with behavior suggests that they may be linked to successful comprehension or calculation.

**Significance Statement:** Neural processing of arithmetic relies on dedicated, modality independent cortical networks that are distinct from those underlying language processing. Using a simultaneous cocktail party listening paradigm, we found that these separate networks segregate naturally when listeners selectively attend to one type over the other. Time-locked activity in the left temporal lobe was observed for responses to both spoken sentences and equations, but the latter additionally showed bilateral parietal activity consistent with arithmetic processing. Critically, these responses were modulated by selective attention and correlated with task behavior, consistent with reflecting high-level processing for speech comprehension or correct calculations. The response dynamics show task-related differences that were used to reliably decode the attentional target of sentences or equations.

## 1. Introduction

Comprehension and manipulation of numbers and words are key aspects of human cognition and share many common features. Numerical operations may rely on language for precise calculations (Spelke and Tsivkin, 2001; Pica et al., 2004) or share logical and syntactic rules with language (Houdé and Tzourio-Mazoyer, 2003). During numerical tasks, frontal, parietal, occipital and temporal areas are activated (Menon et al., 2000; Dehaene et al., 2003, 2004; Arsalidou and Taylor, 2011; Maruyama et al., 2012; Dastjerdi et al., 2013; Harvey et al., 2013; Harvey and Dumoulin, 2017). Bilateral intraparietal sulcus (IPS) is activated by presenting numbers using Arabic or alphabetical notation (Pinel et al., 2001) or speech (Eger et al., 2003). Posterior parietal and prefrontal areas are activated for both arithmetic and language (Price, 2000; Göbel et al., 2001; Venkatraman et al., 2006; Zarnhofer et al., 2012; Bemis and Pylkkänen, 2013). However, some cortical networks activated by numerical stimuli (e.g., IPS), differ from those underlying language processing, even when the stimuli are presented using words (Amalric and Dehaene, 2016, 2019; Monti et al., 2012; Park et al., 2011). Lesion studies (Dehaene and Cohen, 1997; Varley et al., 2005; Baldo and Dronkers, 2007), further provide evidence that the neural basis of numerical processing is distinct from that of language processing (Gelman and Butterworth, 2005; Amalric and Dehaene, 2018).

Beyond cortical location, these neural processes are also dynamic, with dynamic evoked responses to arithmetic arising from parietal, occipital, temporal and frontal regions (Iguchi and Hashimoto, 2000; Kou and Iwaki, 2007; Ku et al., 2010; Jasinski and Coch, 2012; Maruyama et al., 2012; Iijima and Nishitani, 2017). Arithmetic operations can even be decoded from such responses (Pinheiro-Chagas et al., 2019). Speech studies differentiate early auditory evoked components from later components reflecting linguistic and semantic processing in temporal, parietal and frontal regions (Baggio and Hagoort, 2011; Koelsch et al., 2004; Lau et al., 2008; Obleser et al., 2003, 2004). Linear models called Temporal Response Functions (TRFs), which model time-locked responses to continuous speech (Lalor and Foxe, 2010), also reveal dynamical linguistic processing (Brodbeck et al., 2018a, 2018b; Broderick et al., 2018; Di Liberto et al., 2015).

To investigate cortical processing of spoken language and arithmetic, we utilize a technique pioneered by Ding et al. (2016) of presenting isochronous (fixed rate) words and sentences. There, the single syllable word rate, also the dominant acoustic rate, is tracked strongly by auditory neural responses, as expected. However, cortical responses also strongly track the sentence rate, completely absent in the acoustics, possibly reflecting hierarchical language processing (Sheng et al., 2018; Jin et al., 2020; Luo and Ding, 2020). When subjects selectively attend to one speech stream among several, in a ‘cocktail party paradigm’, the sentence rate is tracked only for the attended speaker (Ding et al., 2018). Similarly, cocktail party studies using TRFs show early auditory responses irrespective of attention, and later attention-modulated responses to higher order speech features (Ding and Simon, 2012; Brodbeck et al., 2018b). Attention modulates activation related to numerical processing as well (Castaldi et al., 2019).

Here, magnetoencephalography (MEG) is used to study the cortical processing of short spoken sentences and simple arithmetic equations, presented simultaneously at fixed sentence, equation, word and symbol rates, in an isochronous cocktail party paradigm. This study is motivated by several questions, of increasing complexity. The most basic is whether isochronously presented equations allow segregation of equation-level from symbol-level neural processing in the frequency domain. We demonstrate strong evidence for this segregation. The next level is whether equation- and sentence-level processing show shared or distinct cortical activity areas. We demonstrate evidence for both (shared activity in the left temporal lobe, and distinct equation processing in bilateral IPS and occipital lobe). Finally, we address whether the cocktail party listening paradigm can further differentiate between them, and we find that it does: selective attention allows greater differentiation between the higher-level processing, and, critically, also surfaces neural correlations with behavioral measures.

## 2. Methods

### 2.1. Participants

MEG data was collected from 22 adults (average age 22.6 yrs, 10 female, 21 right handed) who were native English speakers. The participants gave informed consent and received monetary compensation. All experimental procedures were approved by the Internal Review Board of the University of Maryland, College Park. To ensure that the subjects could satisfactorily perform the arithmetic task, only subjects who self-reported that they had taken at least one college level math course were recruited.

### 2.2. Speech stimuli

Monosyllabic words were synthesized with both male and female speakers using the ReadSpeaker synthesizer (https://www.readspeaker.com, ‘James’ and ‘Kate’ voices). The language stimuli consisted of 4-word sentences and the arithmetic stimuli consisted of 5-word equations (hereafter, arithmetic words are referred to as ‘symbols’, arithmetic sentences as ‘equations’, non-arithmetic words as ‘words’ and non-arithmetic sentences as ‘sentences’). The words and symbols were modified to be of constant durations to allow for separate word, symbol, sentence, and equation rates, so that the neural response to each of these could be separated in the frequency domain. The words and symbols were constructed with fixed durations of 375 ms and 360 ms respectively, giving a word rate of 2.67 Hz, a symbol rate of 2.78 Hz, a sentence rate of 0.67 Hz, and an equation rate of 0.55 Hz. All the words and symbols were mono-syllabic, and hence the syllabic rate is identical to the word/symbol rate. These rates are quite fast for spoken English, and, though intelligible, can be difficult to follow in the cocktail party conditions. Because neural signals below 0.5 Hz are very noisy, however, it was not deemed appropriate to reduce the rates further; preliminary testing showed that these rates were a suitable compromise between ease of understanding and reasonable neural signal to noise ratio. In addition, the rates were selected such that 10 equations and 12 sentences (each of which make up a single “trial”) would have the same duration (18 s), allowing for precise frequency resolution at both these rates.

The individual words and symbols were shortened by removing silent portions before their beginning and after their end, and then manipulated to give durations of 375 ms and 360 ms respectively, using the overlap-add resynthesis method in Praat (Boersma and Weenick, 2018). The words and symbols were respectively formed into sentences and equations (described below) and were lowpass filtered below 4 kHz using a 3^rd^ order elliptic filter (the air-tube system used to deliver the stimulus has a lowpass transfer function with cutoff approximately 4 kHz). Finally, each stimulus was normalized to have approximately equal perceptual loudness using the MATLAB ‘integratedLoudness’ function.

The equations were constructed using a limited set of symbols consisting of the word ‘is’ (denoting ‘=’), three operators (‘plus’ (+), ‘less’ (−) and ‘times’ (×)), and the eleven English monosyllabic numbers (‘nil’, ‘one’ through ‘six’, ‘eight’ through ‘ten’, and ‘twelve’). The equations themselves consisted of a pair of monosyllabic operands (numbers) joined by an operator, an ‘is’ statement of equivalence, and a monosyllabic result; the result could be either the first or last symbol in the equation (e.g., ‘three plus two is five’ or ‘five is three plus two’). The equations were randomly generated with repetitions allowed, in order to roughly balance the occurrences of each number (although smaller numbers are still more frequent since there are more mathematically correct equations using only the smallest numbers). The fact that there were a limited set of symbols and that the same symbol ‘is’ occurs in every sentence, in either the 2^nd^ or 4^th^ position, are additional regularities, which contribute to additional peaks in the acoustic stimulus spectrum at the first and second harmonic of the equation rate (1.11 Hz and 1.66 Hz) as seen in Fig. 1 (and borne out by simulations). Although less than ideal, it is difficult to avoid in a paradigm when restricting to mathematically well-formed equations. Hence, we do not analyze the neural responses at those harmonic frequencies, since their relative contributions from auditory vs. arithmetic processing are not simple to estimate. The sentences were also constructed with two related syntactic structures to be similar to the two equation formats: verb in second position (e.g., ‘cats drink warm milk’) and verb in third position (e.g., ‘head chef bakes pie’), but unlike the ‘is’ of the arithmetic case, the verb changed with every sentence and there were no analogous harmonic peaks in the sentence case. Deviants were also constructed: deviant equations were properly structured but mathematically incorrect (e.g., ‘one plus one is ten’); analogously, deviant sentences were syntactically correct but semantically nonsensical (e.g., ‘big boats eat cake’). Cocktail party stimuli were constructed by adding the acoustic waveforms of the sentences and equations in a single audio channel (see Fig. 1) and presented diotically (identical for both ears). The speakers were different (male and female), in order to simplify the task of segregating diotically presented speech. The mixed speech was then normalized to have the same loudness as all the single speaker stimuli using the abovementioned algorithm.

**Figure 1:**
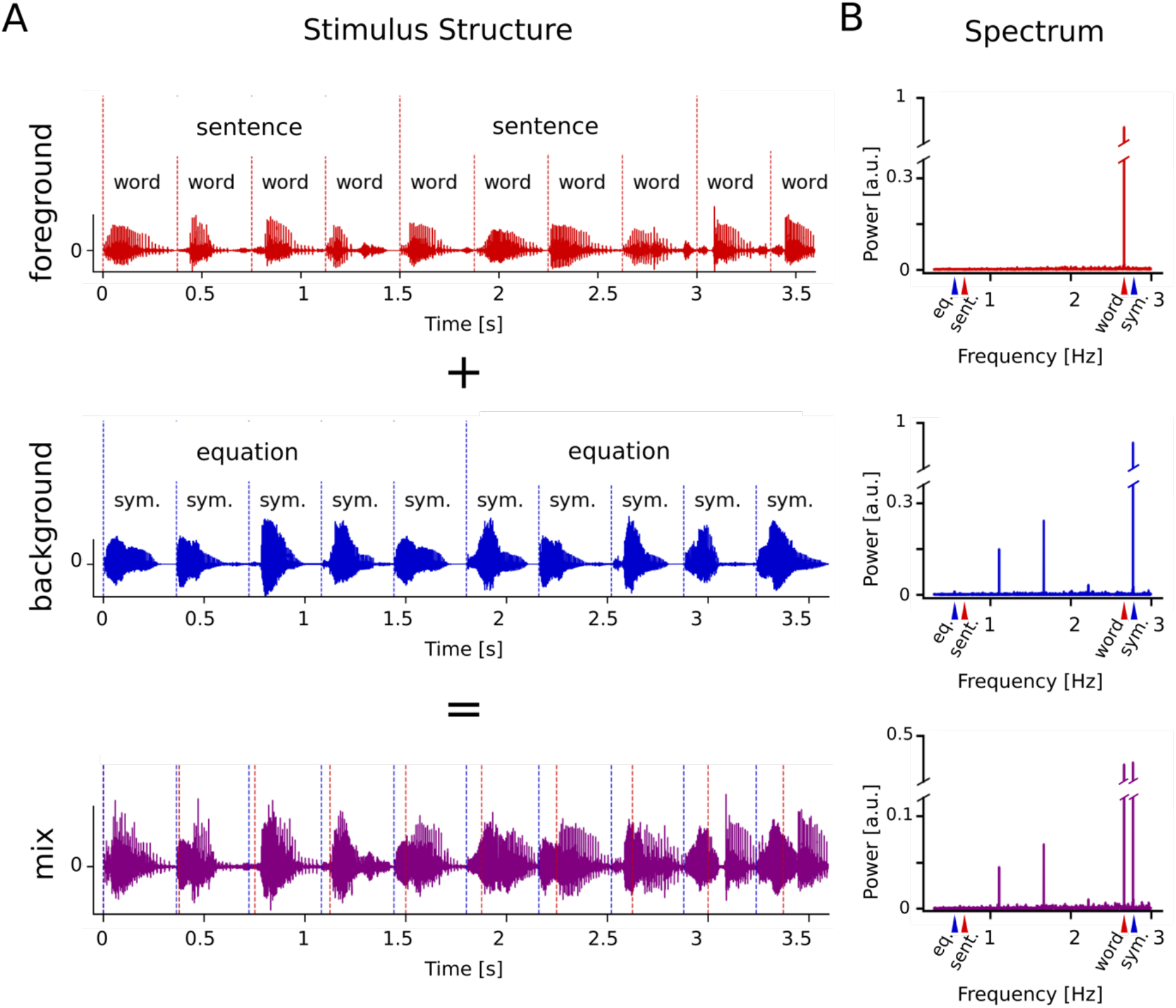
Stimulus structure. **A.** The foreground, background, and mix waveforms for the initial section of the stimulus for a two-speaker attend-language trial. The sentence, equation, word, and symbol structures are shown. The word and symbol rhythms are clearly visible in the waveforms. The mix was presented diotically and is the linear sum of both streams. **B.** The frequency spectrum of the Hilbert envelope of the entire concatenated stimulus for the attend-sentences condition (432 s duration). The sentence (0.67 Hz), equation (0.55 Hz), word (2.67 Hz) and symbol (2.78 Hz) rates are indicated by colored arrows under the x-axis. Clear word and symbol rate peaks are seen in the foreground and background respectively, while the mix spectrum has both peaks. Note that there are no sentence rate or equation rate peaks in the stimulus spectrum. The appearance of harmonics of the equation rate are consistent with the limited set of math symbols used.

### 2.3. Experimental design

The experiment was conducted in blocks: 4 single speaker blocks (2 × 2: male and female, sentences and equations) were followed by 8 cocktail party blocks (see Table 1). The order of the gender of the speaker was counterbalanced across subjects. Each block consisted of multiple trials (10 for single speaker, 6 for cocktail party as shown in Table 1). 50% of the blocks had one deviant trial. Each trial consisted of 10 equations or 12 sentences (or both, for cocktail party conditions) and was 18 s in duration for all cases (0.360 s/symbol × 5 symbols/equations × 10 equations = 18 s; 0.375 s/word × 4 words/sentence × 12 sentences = 18 s). In total, the single speaker conditions had 240 sentences and 200 equations, and the cocktail party conditions had 288 sentences and 240 equations in the foreground. Deviant trials had 4 equations or 5 sentences being deviants. At the start of each block, the subject was instructed which stimulus to attend to, and was asked to press a button at the end of each trial to indicate whether a deviant was detected (right button: yes; left button: no). The subjects kept their eyes open, and a screen indicated which voice they should attend to (‘Attend Male’ or ‘Attend Female’) while the stimulus was presented diotically. After each trial, the stimulus was paused, and the screen displayed the text ‘Outlier?’ until the subjects pressed one of the two buttons. There was a 2-second break after the button press, after which the next trial stimulus was presented.

**Table 1:** Experiment Block Structure. The experiment consisted of 4 single speaker blocks followed by 8 cocktail party blocks. Each trial was 18 s in duration and consisted of 10 equations (1.8 s × 10 = 18 s) or 12 sentences (1.5 s × 12 = 18 s). The speaker gender was counterbalanced across subjects (i.e., the order of column 3 was changed).

| **Table 1: Experiment Block Structure** |  |  |  |
| --- | --- | --- | --- |
| Foreground | Background | Speaker<br>Foreground<br>(Background) | Number of trials<br>per block |
| Equations | - | Male | 10 |
| Sentences | - | Male | 10 |
| Equations | - | Female | 10 |
| Sentences | - | Female | 10 |
| Sentences | Equations | Female (Male) | 6 |
| Equations | Sentences | Female (Male) | 6 |
| Sentences | Equations | Male (Female) | 6 |
| Equations | Sentences | Male (Female) | 6 |
| Sentences | Equations | Female (Male) | 6 |
| Equations | Sentences | Female (Male) | 6 |
| Sentences | Equations | Male (Female) | 6 |
| Equations | Sentences | Male (Female) | 6 |

Since the deviant detection task was challenging, especially in the cocktail party case, subjects were asked to practice detecting deviants just before they were placed inside the MEG scanner (2 trials of language, 2 trials of arithmetic, with stimuli not used during the experiment). The majority of subjects reported that it was easier to follow and detect deviants in the equations compared to the sentences. This might arise for several reasons, e.g., because the equations had a restricted set of simple numbers, or because the repetitive ‘is’ symbol helped keep track of equation structure.

This experiment was not preregistered. The data has been made available at <URL to be provided upon acceptance> and the code is available at <URL to be provided upon acceptance>.

### 2.4. MEG data acquisition and preprocessing

A 157 axial gradiometer whole head MEG system (Kanazawa Institute of Technology, Nonoichi, Ishikawa, Japan) was used to record MEG data while subjects rested in the supine position in a magnetically shielded room (VAC, Hanau, Germany). The data was recorded at a sampling rate of 2 kHz with an online 500 Hz low pass filter, and a 60 Hz notch filter. Saturating channels were excluded (approximately two channels on average) and the data was denoised using time-shift principal component analysis (de Cheveigné and Simon, 2007) to remove external noise, and sensor noise suppression (de Cheveigné and Simon, 2008) to suppress channel artifacts. All subsequent analyses were performed in mne-python 0.19.2 (Gramfort, 2013; Gramfort et al., 2014) and eelbrain 0.33 (Brodbeck et al., 2020). The MEG data was filtered from 0.3–40 Hz using an FIR filter (mne-python 0.19.2 default settings), downsampled to 200 Hz, and independent component analysis was used to remove artifacts such as eye blinks, heartbeats, and muscle movements.

### 2.5. Frequency domain analysis

The complex-valued spectrum of the MEG response for each sensor was computed using the Discrete Fourier Transform (DFT). The preprocessed MEG responses were separated into 4 conditions: attending math or language, in single speaker or cocktail party conditions. The male and female speaker blocks were combined for all analysis. Within each condition, the MEG responses for each trial were concatenated to form signals of duration 6 minutes for each of the single speaker conditions and 7.2 minutes for each of the cocktail party conditions. The DFT was computed for each sensor in this concatenated response, leading to a frequency resolution of 2.7×10^−3^ Hz for the single speaker conditions and 2.3×10^−3^ Hz for the cocktail party conditions. The amplitudes of the frequency spectra were averaged over all sensors and tested for significant frequency peaks (described in section 2.9).

Frequencies of interest were selected corresponding to the equation rate (0.555 Hz), the sentence rate (0.667 Hz), the symbol rate (2.778 Hz), and the word rate (2.667 Hz). Note that the duration of the signals is an exact multiple of both the symbol and the word durations, ensuring that the frequency spectrum contained an exact DFT value at each of these four rates. In addition, the neighboring 5 frequency values (width of ∼0.01 Hz) on either side of these key frequencies were also selected to be used in a noise model for statistical tests.

### 2.6. Neural source localization

The head shape of each subject was digitized using a Polhemus 3SPACE FASTRAK system, and head position was measured before and after the experiment using five marker coils. The marker coil locations and the digitized head shape were used to co-register the template FreeSurfer ‘fsaverage’ brain (Fischl, 2012) using rotation, translation and uniform scaling. A volume source space was formed by dividing the brain volume into a grid of 12 mm sized voxels. This source space was used to compute an inverse operator using minimum norm estimation (MNE) (Hämäläinen and Ilmoniemi, 1994), with a noise covariance estimated from empty room data. The complex-valued sensor distributions (with both amplitude and phase) of each of the 44 selected frequencies (4 frequencies of interest with 10 sidebands each) are given by the Fourier transform of each sensor’s responses (concatenated over trials) at that frequency, in each condition. Each complex-valued sensor distribution was source-localized independently using MNE onto the volume source space, giving complex-valued source activations (Simon and Wang, 2005). The amplitudes of these complex-valued source activations were used for subsequent analysis. The sideband source distributions were averaged together to form the noise model.

### 2.7. Temporal Response Functions (TRFs)

The preprocessed single-trial MEG responses in each of the four conditions (excluding deviant trials) were source-localized in the time domain using MNE, similar to the method described above in the frequency domain. The MEG signals were further lowpassed below 10 Hz using an FIR filter (default settings in mne python) and downsampled to 100 Hz for the TRF analysis. These responses were then used along with representations of the stimulus to estimate TRFs. The linear TRF model for *P* predictors (stimulus representations) is given by

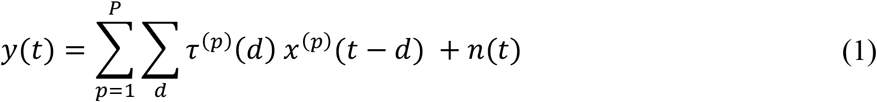

where *y*(*t*) is the response at a neural source at time *t*, *x*^(*p*)^(*t* − *d*) is the time shifted *p*^th^ predictor (e.g., speech envelope, word onsets, etc., as explained below) with time lag of *d*, *τ*^(*p*)^(*d*) is the value of the TRF corresponding to the *p*^th^ predictor at lag *d*, and *n*(*t*) is the residual noise. The TRF estimates the impulse response of the neural system for that predictor, and can be interpreted as the average time-locked response to continuous stimuli (Lalor and Foxe, 2010). For this analysis, several predictors were used to estimate TRFs at each neural source using the boosting algorithm (David et al., 2007), as implemented in eelbrain, thereby separating the neural response to different features. The boosting algorithm may result in overly sparse TRFs, and hence an overlapping basis of 30 ms Hamming windows (with 10 ms spacing) was used in order to allow smoothly varying responses. For the volume source space, the TRF at each voxel for a particular predictor is a vector that varies over the time lags, representing the amplitude and direction of the current dipole activity.

The stimulus was transformed into 2 types of representations that were used for TRF analysis: acoustic envelopes and rhythmic word/symbol or sentence/equation onsets. Although we were primarily interested in responses to sentences and equations, a linear model with only sentence/equation onsets would be disadvantaged by the fact that these representations are highly correlated with the acoustics. Hence by jointly estimating the acoustic envelope and word onset TRFs in the model, the lower-level acoustic responses are automatically separated, allowing the dominantly higher-level processing to emerge in the sentence/equation TRFs. The acoustic envelope was constructed using the 1-40 Hz bandpassed Hilbert envelope of the audio signal (FIR filter used above). The onset representations were formed by placing impulses at the regular intervals corresponding to the onset of the corresponding linguistic unit. The four onset responses were: impulses at 375 ms spacing for word onsets, 360 ms for symbol onsets, 1500 ms for sentence onsets, and 1800 ms for equation onsets. Values at all other time points in these onset representations were set to zero. In order to separate out responses to stimulus onset and offset, the first and last sentences were assigned separate onset predictors, which were not analyzed further except to note that their TRFs showed strong and sustained onset and offset responses, respectively. The remaining (middle) sentences’ onsets were combined into one predictor that was used for further TRF analysis. The same procedure was followed for the equation onset predictors.

For each of the two single-speaker conditions, 5 predictors were used in the TRF model: the corresponding three sentence/equation onsets (just described), word/symbol onsets and the acoustic envelope. For each of the two cocktail conditions, 10 predictors were used in the TRF model: the abovementioned 5 predictors, for each of the foreground and the background stimuli. The predictors were fit in the TRF model jointly, without any preference given to one of them over another.

The TRF for the speech envelope and the word/symbol onsets were estimated for time lags of 0-350 ms in order to limit the TRF duration to before the onset of the next word (at 375 ms) or symbol (at 360 ms). The sentence and equation TRFs were estimated starting from 350 ms to avoid onset responses, as well as lagged responses to the previous sentence. The sentence TRF was estimated until 1850 ms (350 ms past the end of the sentence) and the equation TRF was estimated until 2150 ms (350 ms past the end of the equation), in order to detect lagged responses. These sentence and equation TRFs were used to further analyze high level arithmetic and language processing.

### 2.8. Decoder analysis

All decoding analyses were performed using scikit-learn (Pedregosa et al., 2011) and mne-python software. To investigate the temporal dynamics of responses, linear classifiers were trained on the MEG sensor space signals bandpassed 0.3-10 Hz at 200 Hz sampling frequency. Decoders were trained directly on the sensor space signals, since the linear transformation to source space cannot increase the information already present in the MEG sensor signals. The matrix of observations **x** ∈ ℝ*^N^*^×*M*^, for *N* samples and *M* sensors in each sample, was used to predict the vector of labels **y** ∈ {0, 1}*^N^* at each time point of sentences or equations. The labels correspond to the two attention conditions (attend-equations or attend-sentences). The decoders were trained in the single speaker conditions on time points from 0 to 1500 ms for both sentences (duration 1500 ms) and equations (duration 1800 ms). Therefore, the decoder at each time point learns to predict the attended stimulus type (equations or sentences) using the MEG sensor topography at that time point.

In a similar manner, the operator type in the arithmetic condition was also decoded from the MEG sensor topographies at each time point, in the 720 ms time window of each equation that contained the operator and its subsequent operand. 3 decoders were trained for the 3 comparisons (‘plus’ vs. ‘less’, ‘less’ vs. ‘times’ and ‘plus’ vs. ‘times’).

To further investigate the patterns of cortical activity, linear classifiers were trained on the source localized MEG responses at each voxel, with **x** ∈ ℝ*^N^*^×*T*^ for *N* samples and *T* time points in each sample. The response dynamics of the entire sentence/equation may not be suitable for decoding: since the equations are comprised of five symbols, while the sentences of four words, this might lead to decoding the equations vs. sentences based on whether there were five vs. four auditory responses to acoustic onsets. To minimize this confound, two types of classifiers were used based on responses to only one word/symbol (and hence with only one acoustic onset). 1) Decoding based on first words: The first symbol of each equation and first word of each sentence was used as the sample, with a label denoting attend equations or attend sentences conditions. 2) Decoding based on last words: The last symbol or word was used. Words of duration 375 ms were downsampled to match the duration of the symbols (360 ms), in order to have equal length training samples. This method was used separately for both the single speaker and the cocktail party conditions. The decoder at each voxel learns to predict the attended stimulus type (equations or sentences) using the temporal dynamics of the response at that voxel.

Finally, the effect of attention was investigated using two sets of classifiers for equations and sentences at each voxel. For the attend-equations classifier, the cocktail party trials were separated into samples at the equation boundaries (12 equations per trial), and the labels were marked as ‘1’ when math was attended to and ‘0’ when not. The time duration T was 0-1800 ms (entire equation). For the attend-sentences classifier, the cocktail party trials were separated into samples at the sentence boundaries (10 sentences per trial) and the labels were ‘1’ when attending to sentences and ‘0’ otherwise. The time duration T was 0-1500 ms (entire sentence). Therefore, the attend-equations decoder at each voxel learns to predict whether the equation stimulus was attended to using the temporal dynamics of the response to the equation at that voxel (and similarly for the attend-sentences decoder).

In summary, the decoders at each time point reveal the dynamics of decoding attention to equations vs. sentences from MEG sensor topographies, and the decoders at each voxel reveal the ability to decode arithmetic and language processing in specific cortical areas. The trained classifiers were tested on a separate set and the score of the decoder was computed. Logistic regression classifiers were used, with 5-fold cross-validation, within-subject for all the trials. The area under the receiver operating characteristic curve (*AUC*) was used to quantify the performance of the classifiers.

### 2.9. Statistical analysis

The amplitude spectrum for each condition was averaged across sensors, and permutation tests were used to detect significant peaks across subjects (n=22). Each frequency value in the spectrum from 0.3 to 3 Hz was tested for a significant increase over the neighboring 5 values on either side using 10000 permutations and the max-t method (Nichols and Holmes, 2002) to adjust for multiple comparisons. For these tests, we report the p-values and the t-values, and deliberately omit the degrees of freedom to avoid direct comparison between the two, since the p-values are derived entirely from the permutation distribution and not from the t-distribution. Correlation tests were also performed to investigate associations between different responses (e.g., sentence rate vs. equation rate) within each subject. Pearson correlation tests with Holm-Bonferroni correction were used on the responses at the frequencies of interest, after subtracting the average of the five neighboring bins on either side.

The source distributions for each individual were mapped onto the FreeSurfer ‘fsaverage’ brain, in order to facilitate group statistics. To account for individual variability and mislocalization during this mapping, the distributions were spatially smoothed using a Gaussian window with a standard deviation of 12 mm for all statistical tests. The source localized frequency responses were tested for a significant increase over the corresponding noise model formed by averaging the source localized responses of the five neighboring frequencies on either side. Nonparametric permutation tests (Nichols and Holmes, 2002) and Threshold Free Cluster Enhancement (TFCE) (Smith and Nichols, 2009) were performed to compare the response against the noise and to control for multiple comparisons. A detailed explanation of this method can be found in Brodbeck et al. (2018). Briefly, a test statistic (in this case, paired samples t-statistics between true responses and noise models) is computed for the true data and 10000 random permutations of the data labels. The TFCE algorithm is applied to these statistics, in order to enhance continuous clusters of large values and a distribution consisting of the maximum TFCE value for each permutation is formed. Any value in the original TFCE map that exceeds the 95th percentile is considered significant at the 5% significance level. In all subsequent results, the minimum p-value and the maximum or minimum t-value across voxels is reported as *p_min_*, *t_max_* or *t_min_* respectively. Note that the *p_min_* is derived from the permutation distribution and cannot be derived directly from *t_max_* or *t_min_* using the t-distribution (degrees of freedom are also omitted due to this reason). Lateralization tests were performed by testing each voxel in the left hemisphere with the corresponding voxel in the right, using permutation tests and TFCE with paired samples t-statistics. For the attend math conditions, equations were separated by operator type (‘plus’ (+), ‘less’ (−) or ‘times’ (×)) to test for specific responses to each operator. No significant differences were found between the source localized responses to each operator.

To test for significant effects and interactions, repeated measures ANOVAs were performed on the source localized responses at each frequency of interest after subtracting the corresponding noise model. Nonparametric permutation tests with TFCE were used to correct for multiple comparisons, similar to the method described above. In brief, a repeated-measures ANOVA is performed at each voxel, and then, for each effect or interaction, the voxel-wise F-values from this ANOVA are passed into the TFCE algorithm, followed by permutation tests as described earlier. This method detects significant clusters in source space for each significant effect. Note that the maximum F-value in the original map within a cluster (*F_max_*) and the p-value of the cluster are reported (and degrees of freedom omitted), for the same reasons as those explained in the previous paragraph (i.e., p-values are derived from the permutation distribution and not the F-distribution).

Several types of repeated measures ANOVAs were performed using the abovementioned method. In the single speaker case, a 2 × 2 ANOVA with factors stimulus (‘language’ for words/sentences or ‘math’ for symbols/equations) and frequency (low for sentence/equation and high for word/symbol) was performed. For the cocktail party case, a 2 × 2 × 2 ANOVA with the added factor of attention (attended or unattended) was performed. In addition, two further ANOVAs were performed to investigate hemispheric effects, using an additional factor of hemisphere for both the single speaker (2 × 2 × 2 ANOVA) and the cocktail party (2 × 2 × 2 × 2 ANOVA) conditions.

To investigate significant ANOVA effects further, post-hoc t-tests across subjects were performed on the responses averaged across voxels within the relevant significant cluster. For this scalar t-test, a Holm-Bonferroni correction was applied to correct for multiple comparisons. For these tests, the t-values with degrees of freedom, corrected p-values and Cohen’s d effect sizes are reported.

Behavioral responses for the deviant detection task were classified as either correct or incorrect, and the number of correct responses for each subject was correlated with the source localized response power of that subject. The noise model for each frequency of interest was subtracted from the response power before correlating with behavior. Nonparametric permutation tests with TFCE were used in a manner similar to that given above. The only difference was that the statistic used for comparison was the Pearson correlation coefficient between the two variables (behavior and response power), and the maximum correlation coefficient across voxels is reported as *r_max_*.

The TRFs were tested for significance using vector tests based on Hotelling’s T^2^ statistic (Mardia, 1975). Since the TRFs consist of time varying vectors, this method tests for consistent vector directions across all subjects at each time point and each voxel. The Hotelling’s T^2^ statistic was used with non-parametric permutation tests and TFCE as described above, with the added dimension of time, and the maximum T^2^ statistic across voxels is reported as 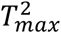. This statistic is more suitable than a t-statistic based on the amplitude of the TRF vectors, since activation from distinct neural processes may have overlapping localizations (due to the limited resolution of MEG), but different current directions. Similarly, for the cocktail party conditions, foreground and background TRFs were compared in order to assess attentional effects, using the difference vector of the two TRFs.

Finally, the decoders were tested across subjects for a significant increase in decoding ability above chance (*AUC* = 0.5) at each time point or at each voxel. Multiple comparisons were controlled for using permutation tests and TFCE, similar to the above cases, with *AUC* as the test statistic.

## 3. Results

### 3.1 Behavioral results

After each trial, subjects indicated whether that trial contained a deviant by pressing a button. The single speaker conditions had higher deviant detection accuracies (equations: mean = 89.9%, std. = 10.7%; sentences: mean = 74%, std. = 13.7%) than the cocktail party conditions (equations: mean = 79.8%, std. = 13.6%; sentences: mean = 62.3%, std = 18.9%). Subjects reported that the equations were perceptually easier to follow than the sentences, consistent with the fact that the equations were formed using a smaller set of symbols (restricted to being monosyllabic numbers to preserve the symbol rates). The presence of ‘is’ in each equation may have also contributed to subjects tracking equation boundaries.

### 3.2. Frequency domain analysis

The response power spectrum was averaged over all sensors and a permutation test with the max-t method was performed to check whether the power at each frequency of interest was significantly larger than the average of the neighboring 5 frequency bins on either side (see Fig. 2 A, B) across subjects (n = 22). For the language single speaker condition, the sentence rate (0.67 Hz, *t* = 7.25, *p* < 0.001, Note: degrees of freedom not shown since p-values are derived from the permutation test, see Methods 2.9), its first harmonic (1.33 Hz, *t* = 6.11, *p* = 0.0023), and the word rate (2.67 Hz, *t* = 12.98, *p* < 0.001) were significant (one tailed permutation test of difference of amplitudes with max-t method). Similarly, for the math single speaker condition, the symbol rate (2.78 Hz, *t* = 12.39, *p* < 0.001) and the equation rate (0.55 Hz, *t* = 6.29, *p* = 0.0017) were significant. In this condition, the 1^st^ and 2^nd^ harmonics of the equation rate were also significant (*t* = 7.28, *p* < 0.001 at 1.11Hz; *t* = 7.77, *p* < 0.001 at 1.67 Hz). Thus, in both conditions, the responses track the corresponding sentence or equation rhythms that are not explicitly present in the acoustic signal. The harmonic peak (1.33 Hz) in the language condition is consistent with phrase tracking (Ding et al., 2016), and the harmonics in the arithmetic condition (1.11 Hz, 1.66 Hz) are consistent with auditory processing of acoustic properties of the stimulus associated with the limited number of mathematical symbols employed (see Methods 2.2), or higher-order processing, or both. Correlation tests within subjects, with Holm-Bonferroni correction, were performed on relevant pairs of responses (after subtracting the neighboring 5 bins). Sentence rate responses were significantly correlated with equation rate responses (Pearson’s *r* = 0.576, *p* = 0.015). Word rate responses were significantly correlated with symbol rate responses (*r* = 0.681, *p* = 0.001). Since such correlations may arise from fluctuating degree of task engagement, or variable neural signal to noise ratio across subjects, they were not analyzed further. There were no significant sentence vs. word (*r* = 0.067, *p* > 0.99) or equation vs. symbol (*r* = 0.001, *p* > 0.99) response correlations.

**Figure 2.**
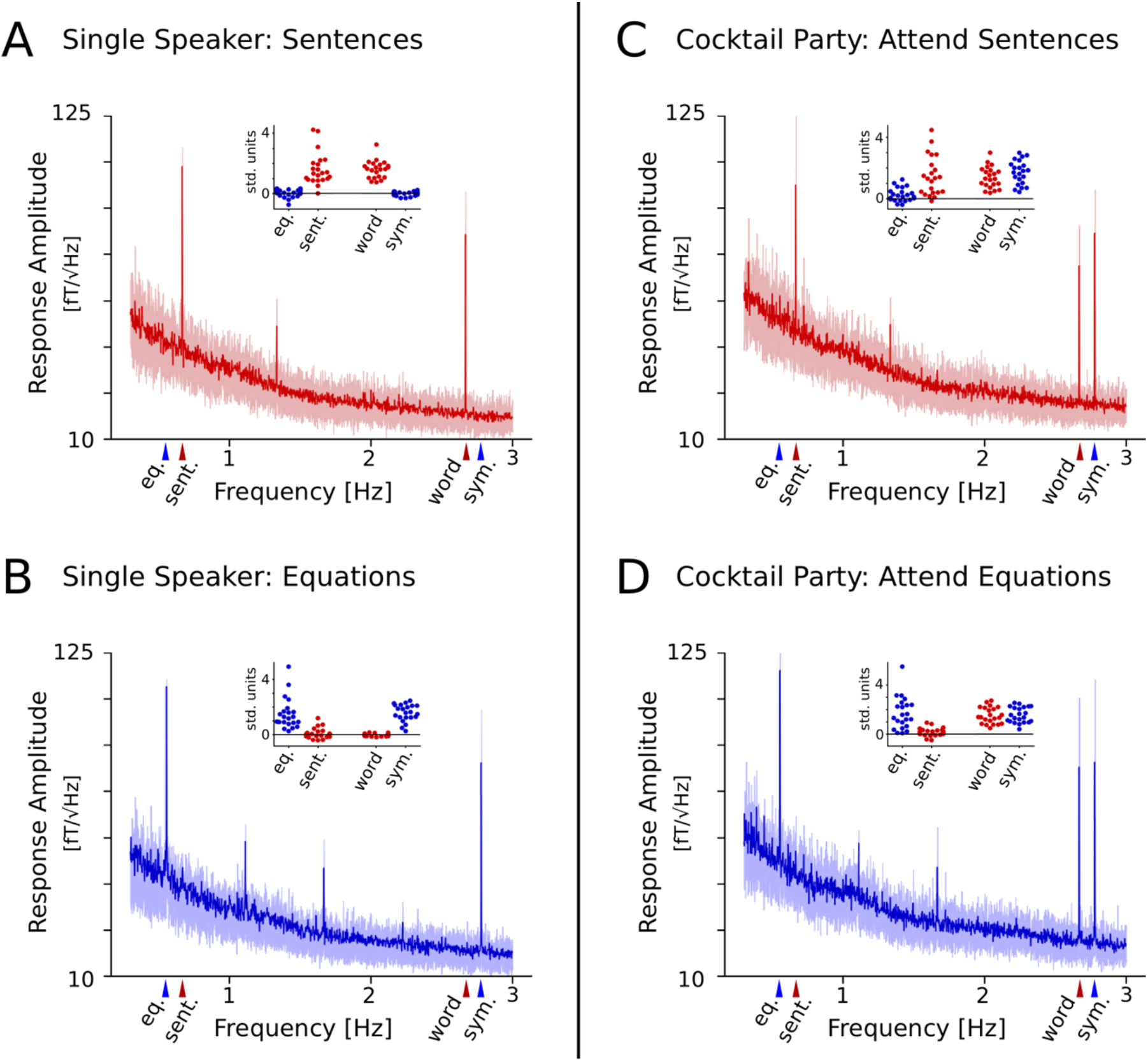
Neural response spectrum. The MEG response spectrum as a function of frequency for the four conditions. The amplitude spectrum, averaged over sensors and subjects, is shown with light shaded regions denoting the 1st-3rd quartile range across subjects. Clear peaks are seen at the sentence, equation, word, and symbol rates (indicated by the arrows under the x-axis). These responses were compared against neighboring bins (of width ∼0.01 Hz, not visible here) for statistical tests. Insets show the average responses at the four frequencies of interest for each subject, after subtracting the neighboring bins. The scale for the insets is standardized within each condition, but with 0 indicating the baseline average activity of the neighboring bins. For the single speaker conditions, peaks appear only at the rates corresponding to the presented stimulus. For the cocktail party conditions, peaks appear at the symbol and word rates regardless of attention, while sentence and equation peaks only appear during the attended condition. There are no analogous sentence or equation peaks during the opposite attention condition.

For the attend-sentences cocktail party condition, only the (attended) word, (unattended) symbol and (attended) sentence rate responses were significant (one tailed permutation test of difference of amplitudes with max-t method; *t* = 9.29, *p* < 0.001; *t* = 10.59, *p* < 0.001; *t* = 5.46, *p* = 0.0176 respectively) as shown in Fig 2C, D. The (unattended) equation rate response was not significant (*t* = 2.99, p > 0.99). On the other hand, for the attend-equations cocktail party condition, the (unattended) word, (attended) symbol and (attended) equation rate responses were significant (*t* = 10.86, *p* < 0.001; *t* = 11.64, *p* < 0.001; *t* = 6.07, *p* = 0.005 respectively), while the (unattended) sentence rate response was not significant (*t* = 2.73 p > 0.99). Responses at the 1^st^ and 2^nd^ harmonics of the equation rate were also significant in the attend-equations condition (1.11 Hz, *t* = 5.31, *p* = 0.027; 1.67 Hz, *t* = 5.09, *p* = 0.04). Correlation tests within subjects were performed, similar to the single speaker case, on all responses except the non-significant unattended sentence and equation rates. Once again, attended sentence rate responses were significantly correlated with attended equation rate responses (*r* = 0.68, *p* = 0.0023). Word rate responses were significantly correlated with symbol rate responses for both attended (*r* = 0.69, *p* = 0.0021) and unattended cases (*r* = 0.83, *p* < 0.001). Other correlations were not significant (attended sentence vs. attended word: *r* = 0.12, *p >* 0.99; attended sentence vs. unattended word: *r* = 0.07, *p* = 0.74; attended equation vs. attended symbol: *r* = 0.49, *p* = 0.083; attended equation vs. unattended symbol: *r* = 0.33, *p* = 0.4).

Since the word and symbol rates are present in the acoustics for both conditions, the neural responses at these rates could merely reflect acoustic processing. However, the fact that the sentence and equation rates are significant only in the corresponding attention condition suggests that these responses may dominantly reflect attention-selective high-level processes. This agrees with prior studies showing a similar effect for language (Ding et al., 2018). Here we show that this effect occurs even for arithmetic equations. However, arithmetic equations are also sentences, so it is unclear from this result alone if the equation peak reflects merely tracking of sentential structure and not arithmetic processing. To investigate this, we used volume source localization on the responses at the relevant frequencies to determine the cortical distribution of these responses.

The responses at the 4 frequencies of interest (word, symbol, sentence and equation rates) were source-localized using the Fourier transform sensor topographies at these frequencies (see Methods 2.6). The amplitudes of the resulting complex-valued volume source distributions were used for all subsequent analysis. For each frequency of interest, the neighboring 5 bins on either side were also source localized (using the same source model) and averaged together to form an estimate of the background noise. The response distributions for each of these frequencies were tested for a significant increase over the noise estimate using nonparametric permutation tests with paired sample t-statistics and TFCE. For both single speaker conditions, the corresponding word or symbol responses were significant (*t_max_* = 12.85, *p_min_* < 0.001, and *t_max_* = 12.77, *p_min_* < 0.001, respectively) in the regions shown in Fig. 3A, B, with the average response being strongest in bilateral auditory cortex. The word and symbol rate responses were not significantly different (*t_min_* = −2.11, *t_max_* = 3.63, *p* > 0.08), consistent with low level auditory processing. The corresponding sentence or equation responses were also significant (*t_max_* = 9.92, *p_min_* < 0.001, and *t_max_* = 7.68, *p_min_* < 0.001, respectively). The source distribution for sentence responses was predominantly in left auditory cortex and temporal lobe, whereas for equations the response was distributed over areas of bilateral temporal, parietal, and occipital lobes. Despite these visually distinct patterns, the two responses were not significantly different (*t_min_* = −3.00, *t_max_* = 3.36, *p* > 0.12), perhaps because large portions of the brain show activity synchronized to the rhythm. Both sentence and equation responses were significantly left lateralized in temporal (*t_max_* = 6.69, *p_min_* < 0.001) and parietal (*t_max_* = 3.9, *p_min_* = 0.009) areas respectively. No significant differences were seen in the responses at the equation rate when separated according to operator type (’+’ vs. ‘-’: *t_min_* = −2.98, *t_max_* = 1.64, *p* > 0.34; ‘-’ vs. ‘×’: *t_min_* = −2.08, *t_max_* = 3.21, *p* > 0.39; ‘×’ vs. ‘+’: *t_min_* = −2.26, *t_max_* = 3.01, *p* > 0.31).

**Figure 3.**
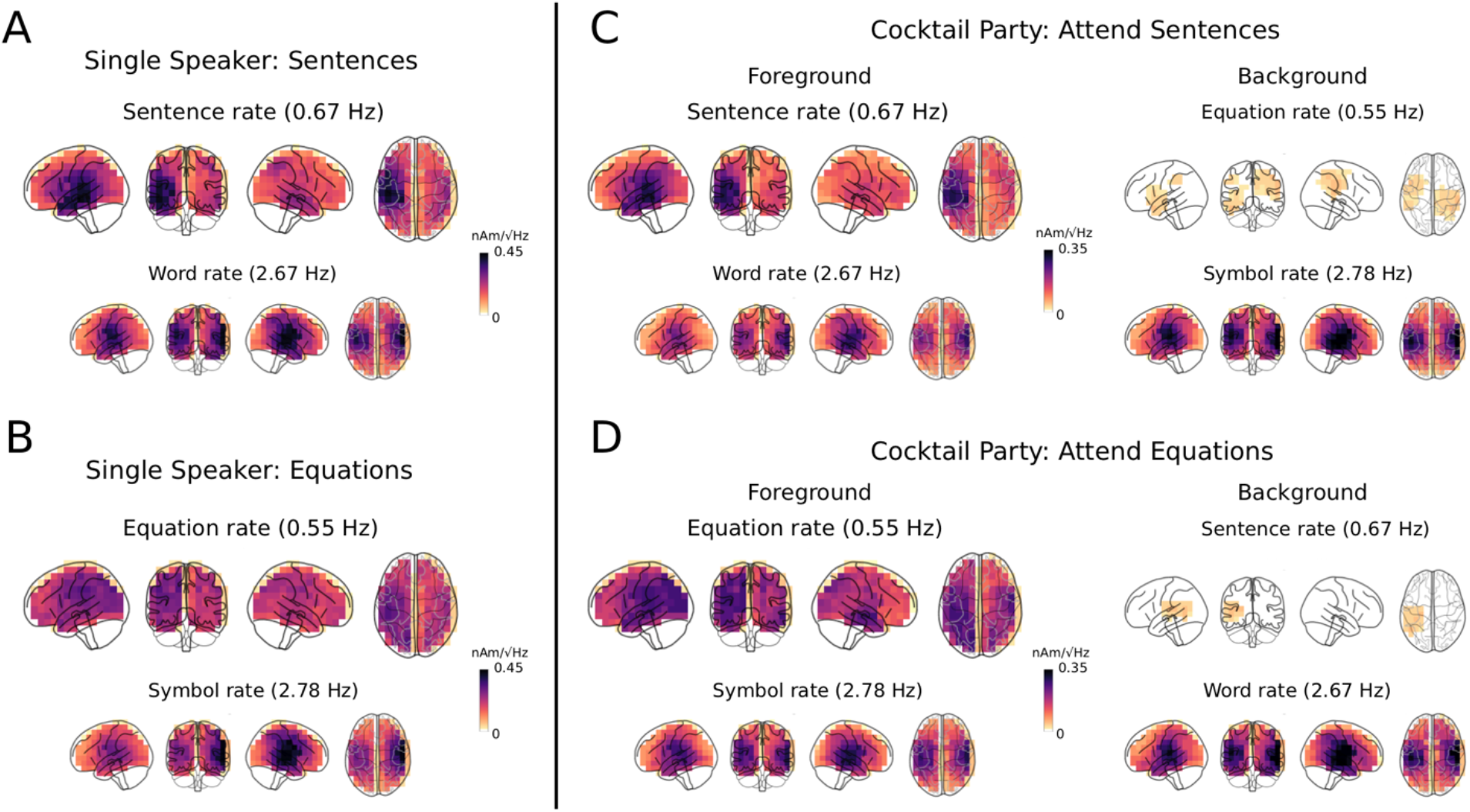
Source localized responses at each frequency of interest. The source localized responses at critical frequencies, averaged over subjects and masked by significant increase over the noise model, are shown. Color scales are normalized within each condition in order to more clearly show the spatial patterns. The word and symbol rate responses are maximal in bilateral auditory cortical areas, while the sentence rate response is maximal in the left temporal lobe. The equation rate responses localize to bilateral parietal, temporal, and occipital areas, albeit with increased left hemispheric activity. Although the background sentence and equation rates also show significant activity, the amplitude of these responses are much smaller than the responses at the corresponding attended rates.

For the cocktail party conditions, similar results were obtained for both word and symbol rate responses (attend sentences: word rate: *t_max_* = 11.9, *p_min_* < 0.001, symbol rate: *t_max_* = 12.8, *p_min_* < 0.001; attend equations: word rate: *t_max_* = 11.1, *p_min_* < 0.001, symbol rate: *t_max_* = 11.01, *p_min_* < 0.001). The response was predominantly in bilateral auditory cortices as shown in Fig. 3C, D, and the symbol and word rates were not significantly different (*t_min_* = −4.31, *t_max_* = 2.33, *p* > 0.16). The attended sentence or equation rate responses were significant (*t_max_* = 6.78, *p_min_* < 0.001, and *t_max_* = 7.87, *p_min_* < 0.001, respectively) and the localization was similar to the single speaker case, albeit more bilateral for the equation rate response. Indeed, the sentence rate response was significantly left lateralized (*t_max_* = 5.36, *p_min_* < 0.001), similar to the single speaker case, but the equation rate response was not (*t_max_* = 2.97, *p* > 0.067). However, the spatial distribution of the equation rate response was larger in the left hemisphere (see Fig. 3); indeed, the source localization of attended sentence responses and attended equation responses were significantly different (*t_min_* = −4.77, *t_max_* = 2.39, *p_min_* = 0.013), with more equation rate responses in the right hemisphere. This indicates that the equation rate response does not originate from the same cortical regions that give rise to the sentence rate response and that the selective attention task is better able to separate these responses. Perhaps surprisingly, the unattended sentence and equation rates were also significant (*t_max_* = 4.02, *p_min_* = 0.005, and *t_max_* = 5.31, *p_min_* < 0.001, respectively) in small clusters, even though such peaks do not appear in the frequency spectrum averaged across all sensors (Fig. 2). Note however, that some individuals did show small peaks at these rates even in the average spectrum (see points above zero for unattended rates in the insets of Fig. 2C, D).

A repeated measures ANOVA was performed for the single speaker case on the abovementioned source space distributions (as shown in Fig. 3) for each frequency of interest. The 2 × 2 ANOVA consisted of factors stimulus (‘language’ for word/sentence or ‘math’ for symbol/equation) and specific frequency (‘high’ for word/symbol or ‘low’ for sentence/equation). The ANOVA was performed on the response at each voxel and cluster-based permutation tests with TFCE were used to correct for multiple comparisons (See Methods 2.9 for choice of reported statistics). The interaction of stimulus × frequency was not significant (*F*_max_ = 10.38, *p* = 0.149, but see below for an interaction effect in an ANOVA with a factor of hemisphere). A significant main effect of frequency (*F*_max_ = 18.63, *p* = 0.006) was found in a right auditory cluster and a significant main effect of stimulus type (*F*_max_ = 21.67, *p* = 0.003) was found in the left auditory/temporal area. Post-hoc t-tests across subjects were performed on the responses averaged across voxels within the significant clusters for each effect; p-values were obtained from the t-distribution and then corrected for multiple comparisons using the Holm-Bonferroni method. These tests revealed that the main effect of stimulus was due to a significant increase in both the sentence over the equation responses (*t*(21) = 2.96, *p* = 0.037, Cohen’s *d* = 0.54) and the word over the symbol responses (*t*(21) = 2.85, *p* = 0.038, Cohen’s *d* = 0.52) in the left auditory/temporal cluster, consistent with increased left temporal activity for language over arithmetic. The main effect of frequency was due to a significant increase in both the word over the sentence responses (*t*(21)=3.67, *p* = 0.01, Cohen’s *d*=1.16), and the symbol over the equation responses (*t*(21)=3.15, *p* = 0.028, Cohen’s *d*=0.97) in the right auditory cluster.

For the cocktail party case, a similar repeated measures ANOVA was performed, but with an additional factor of attention (attended or unattended) leading to a 2 × 2 × 2 design. A significant 3-way interaction of stimulus × attention × frequency was found in a right parietal cluster (*F*_max_ = 15.18, *p* = 0.024). Post-hoc t-tests across subjects with Holm-Bonferroni correction were performed on the responses averaged across voxels within this cluster. These revealed a significant increase in the equation responses compared to the sentence responses when attended (*t*(21) = 3.71, *p* = 0.0103, Cohen’s *d* = 0.82), but no significant difference when unattended (*t*(21) = 2.27, *p* = 0.09, Cohen’s *d* = 0.65). There was also no significant difference between word and symbol responses both when attended (*t*(21)=-0.32, p = 0.75, Cohen’s *d* = −0.06) and unattended (*t*(21)=- 0.69, *p* = 0.99, Cohen’s *d* = −0.09). This is consistent with increased responses to equations in right parietal areas only when attended. In addition to this 3-way interaction, several 2-way interactions and main effects were also detected but were not analyzed further.

Finally, two further ANOVAs were performed with an additional factor of hemisphere for both the single speaker (2 × 2 × 2 ANOVA) and cocktail party (2 × 2 × 2 × 2 ANOVA). For the single speaker case, the 3-way interaction was significant (stimulus × frequency × hemisphere: *F*_max_ = 18.55, *p* = 0.016) in superior parietal voxels. For the cocktail party case, the 4-way interaction was not significant (attention × frequency × stimulus type × hemisphere: *F*_max_ = 8.31, *p*=0.115). However, two 3-way interactions involving hemisphere were significant (attention × frequency × hemisphere: *F*_max_ = 13.71, *p* = 0.031, frequency × stimulus × hemisphere: *F*_max_ = 18.12, *p* =0.017) in temporal voxels. Other effects involving hemisphere were also found to be significant (frequency × hemisphere: *F*_max_ = 41.75, *p* < 0.001, main effect of hemisphere: *F*_max_ = 17.1, *p* = 0.021), as well as several other effects not involving hemisphere. These effects were not analyzed further, but they indicate that the effects of attention, stimulus and frequency depend significantly on the hemisphere, as already suggested by the lateralized clusters found in the simpler ANOVAs described earlier.

In summary, the ANOVA analysis indicates that, in the single speaker case, low-level responses (word/symbol) are significantly stronger than the higher-level responses (sentence/equation) in right auditory areas and that the language responses (sentence/word) are significantly stronger than the arithmetic responses (equation/symbol) in left auditory/temporal areas. Critically, the ANOVA results for cocktail party indicate that the equation responses are significantly larger than the sentence responses in right parietal areas but only when attended to. ANOVAs also indicate that these effects depend on hemisphere as already suggested by the previous pairwise comparisons.

### 3.3. Behavioral correlations

Behavioral performance was correlated with source localized neural responses using non-parametric permutation tests with TFCE, with Pearson correlation as the test statistic. Deviant detection performance for sentences in the single speaker condition was significantly correlated with the sentence rate neural response (*p_min_* = 0.02, maximum correlation in significant regions *r_max_* = 0.62) as shown in Fig. 4. However, detection of equation deviants in the single speaker condition was not significantly correlated with the equation rate neural response; this may be related to the fact that performance in the single speaker arithmetic condition was at ceiling for several participants. The performance when detecting sentence deviants in the cocktail party conditions was correlated with the attended sentence rate response (*r_max_* = 0.62, *p_min_* = 0.015), attended word rate response (*r_max_* = 0.64, *p_min_* = 0.03) as well as the unattended symbol rate response (*r_max_* = 0.79, *p_min_* = 0.001). The performance when detecting equation deviants in the cocktail party condition was correlated with the attended equation rate response (*r_max_* = 0.6, *p_min_* = 0.02) and the unattended word rate response (*r_max_* = 0.74, *p_min_* = 0.04). It was unexpected that the unattended word and symbol rate responses were significantly correlated with behavior, and possible explanations are discussed in section 4.3. Critically, however, sentence and equation rate responses were correlated with behavior only when attended.

**Figure 4.**
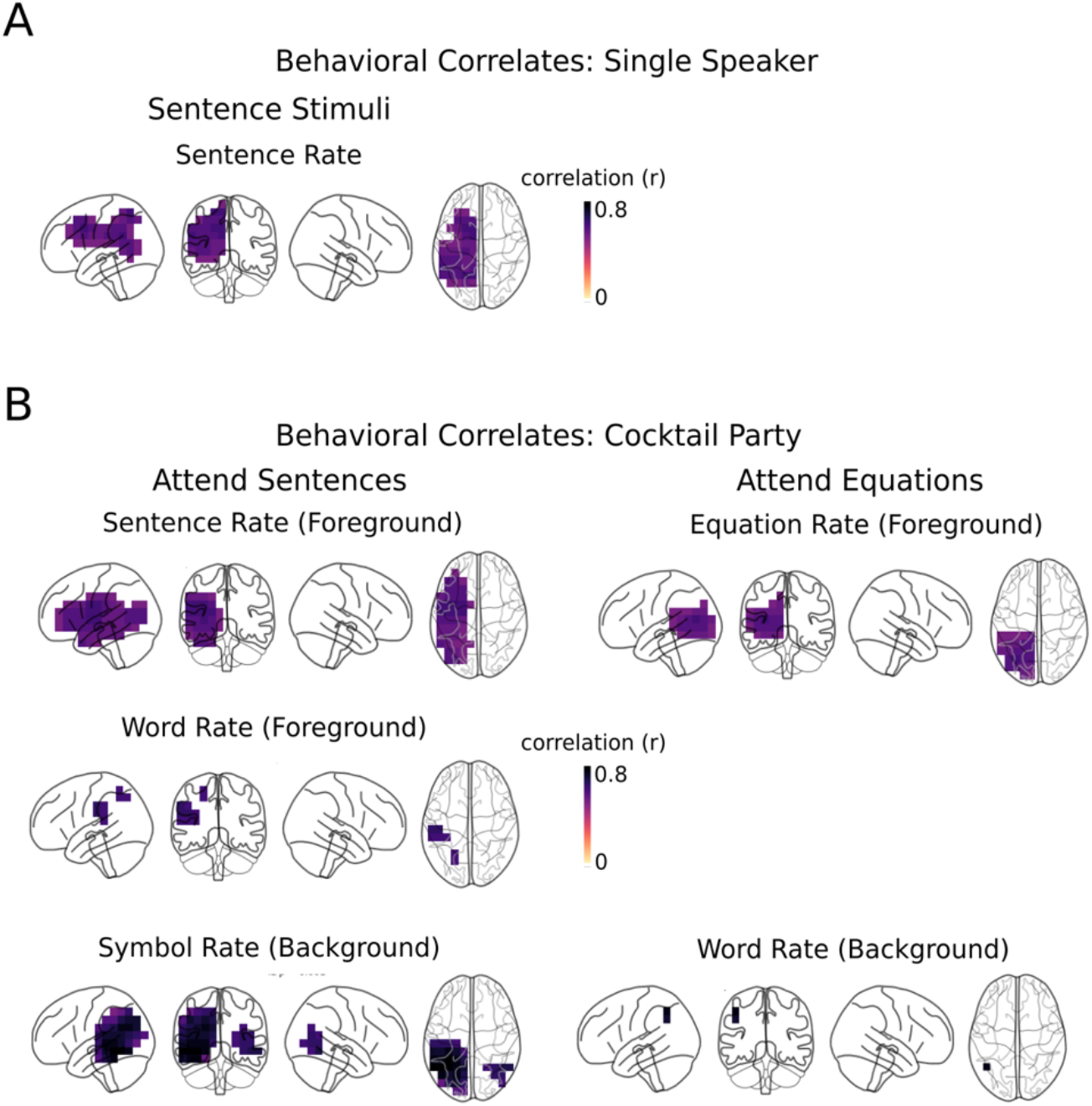
Neural response correlations with behavior. The source localized responses at the frequencies of interest were correlated with the corresponding deviant detection performance, across subjects. The areas of significant correlation are plotted here (same color scale for all plots). Sentence and equation rate responses are significantly correlated with behavior only if attended, while both attended and unattended word rate responses are significantly correlated with behavior. The sentence rate response is significantly correlated over regions in left temporal, parietal, and frontal areas, while significant correlation for the equation rate response is seen in left parietal and occipital regions.

### 3.4. TRF analysis

TRF analysis was performed using source localized MEG time signals for each condition after excluding the deviant trials (details in Methods 2.7). TRFs were simultaneously obtained for responses to the acoustic envelopes, word/symbol onsets and sentence/equation onsets. Although stimuli with fixed and rhythmic word, symbol, sentence, and equation onsets might lend itself to an evoked response analysis, the fact that the words (or symbols) are only separated by 375 ms (or 360 ms) may lead to high-level late responses overlapping with early auditory responses to the next word (or symbol). In contrast, computing simultaneous TRFs to envelopes and word/symbol onsets in the same model as TRFs to equation/sentence onsets regresses out auditory responses from higher-level responses, providing cleaner TRFs for sentences and equations. The obtained envelope and word/symbol TRFs were not used for further analysis, since they were dominated by acoustic responses that have been well-studied in other investigations (Brodbeck et al., 2018a, 2018b). The volume source localized TRFs are time-varying vectors at each voxel. Activity of nearby neural populations may overlap, even if the activity is driven by different processes, due to the limited spatial resolution of MEG. However, these effects may have different current directions due to the anatomy of the cortical surface. Therefore, a test for consistent vector directions, using Hotelling’s T^2^ statistic and permutation tests with TFCE, was used to detect group differences in the direction of current flow (see Methods 2.9).

The sentence and equation TRFs showed significance over several time intervals and many voxels over the duration of the TRF (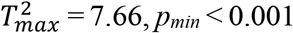, and 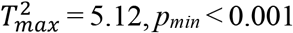, respectively, see Fig 5). The TRFs were computed starting from 350 ms after the sentence onset to 350 ms after the end of the sentence, but because of the fixed-period presentation rate without any breaks between sentences in a trial, the TRFs from 0-350 ms are identical to the last 350 ms. The large peak at the end (and beginning) of each TRF may either arise from processing of the completion of the sentence/equation, or to preparation (or auditory) processing of the new sentence sentence/equation, or both. This peak occurs around 60-180 ms after the start of the new sentence/equation, in the typical latency range of early auditory processing. However, spatial distributions of the peak in the equation TRFs seem to indicate patterns that are not consistent with purely auditory processing, especially for the cocktail party condition (described below). Additionally, significant activity is seen throughout the duration of the sentence/equation that is not tied to word/symbol onsets, indicating that lower-level auditory responses have been successfully regressed out. Therefore, the large peaks at the end plausibly reflect processing of the completion of the sentence/equation (with a latency of 420-530 ms after the last word/symbol). The sentence TRF peaks were significant predominantly in the left temporal lobe, while the equation TRF peaks were significant in bilateral temporal, parietal, and motor areas.

**Figure 5.**
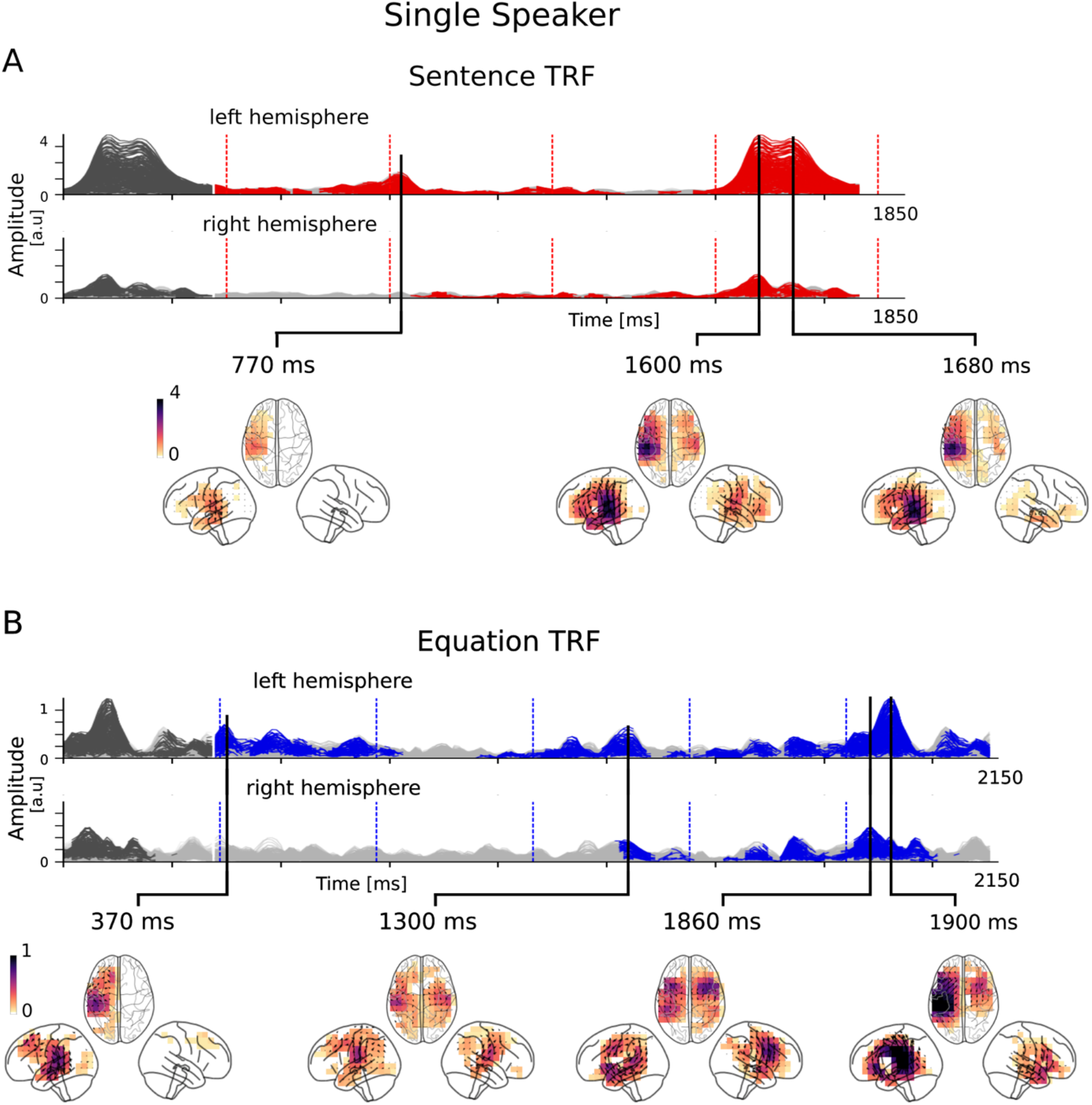
TRFs in the single speaker conditions. Overlay plots of the amplitude of the TRF vectors for each voxel, averaged over subjects. For each TRF subfigure, the top axis shows vector amplitudes of voxels in the left hemisphere and the bottom axis correspondingly in the right hemisphere. Each trace is from the TRF of a single voxel; non-significant time points are shown in gray, while significant time points are shown in red (sentence TRF) or blue (equation TRF). The duration plotted corresponds to that of a sentence or equation, plus 350 ms; because of the fixed presentation rate, the first 350 ms (shown in gray) are identical to the last 350 ms. The large peak at the end (and beginning) of each TRF may either be ascribed to processing of the completion of the sentence/equation, or to the onset of the new sentence sentence/equation, or both. Word and symbol onset times are shown in red and blue dashed lines respectively; it can be seen that response contributions associated with them have been successfully regressed out. Volume source space distributions for several peaks in the TRF amplitudes are shown in the inlay plots, with black arrows denoting current directions (peaks automatically selected as local maxima of the TRFs). Although most of the TRF activity is dominated by neural currents in the left temporal lobe, the equation TRFs show more bilateral activation. The full time-courses of all significant neural source space distributions are shown in Video 1.

#### Video 1. TRFs in the single speaker conditions

The sentence and equation TRFs are shown from 350 ms until 350 ms past the end of the sentence or equation. Traces at significant time points are shown in red (sentence processing) or blue (equation processing). The distribution of cortical activity at each time point is shown below each TRF, with color scales fixed over each entire TRF.

Differences in sentence and equation processing were more readily visible in the TRFs for the cocktail party conditions. The test for consistent vector direction revealed similar results to the single speaker conditions (sentence TRF 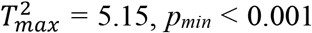, equation 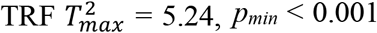) as shown in Fig 6, however, the differences between sentences and equations were more pronounced, especially at for the later peaks 410-600 ms after the onset of the last word or symbol. The peaks in the equation TRF were localized to left motor and parietal regions and right inferior frontal areas that are associated with arithmetic processing. This strengthens the hypothesis that these late peaks indicate lagged higher-level processing of the completed equation and not early auditory/preparatory processing of the subsequent equation. Although the cortical localization of sentence TRF peaks remain consistent in left temporal areas throughout most of the time course, the equation TRF peaks show several distinct cortical patterns, and may reflect distinct processes. The equation TRF showed strong activity in bilateral IPS, superior parietal and motor areas, while sentence TRFs consistently localized predominantly to regions near left auditory cortex (even more so than in the single speaker case). Therefore, selective attention in the cocktail conditions seems to highlight differences between arithmetic and language processing, and possible explanations are discussed section 4.6.

**Figure 6.**
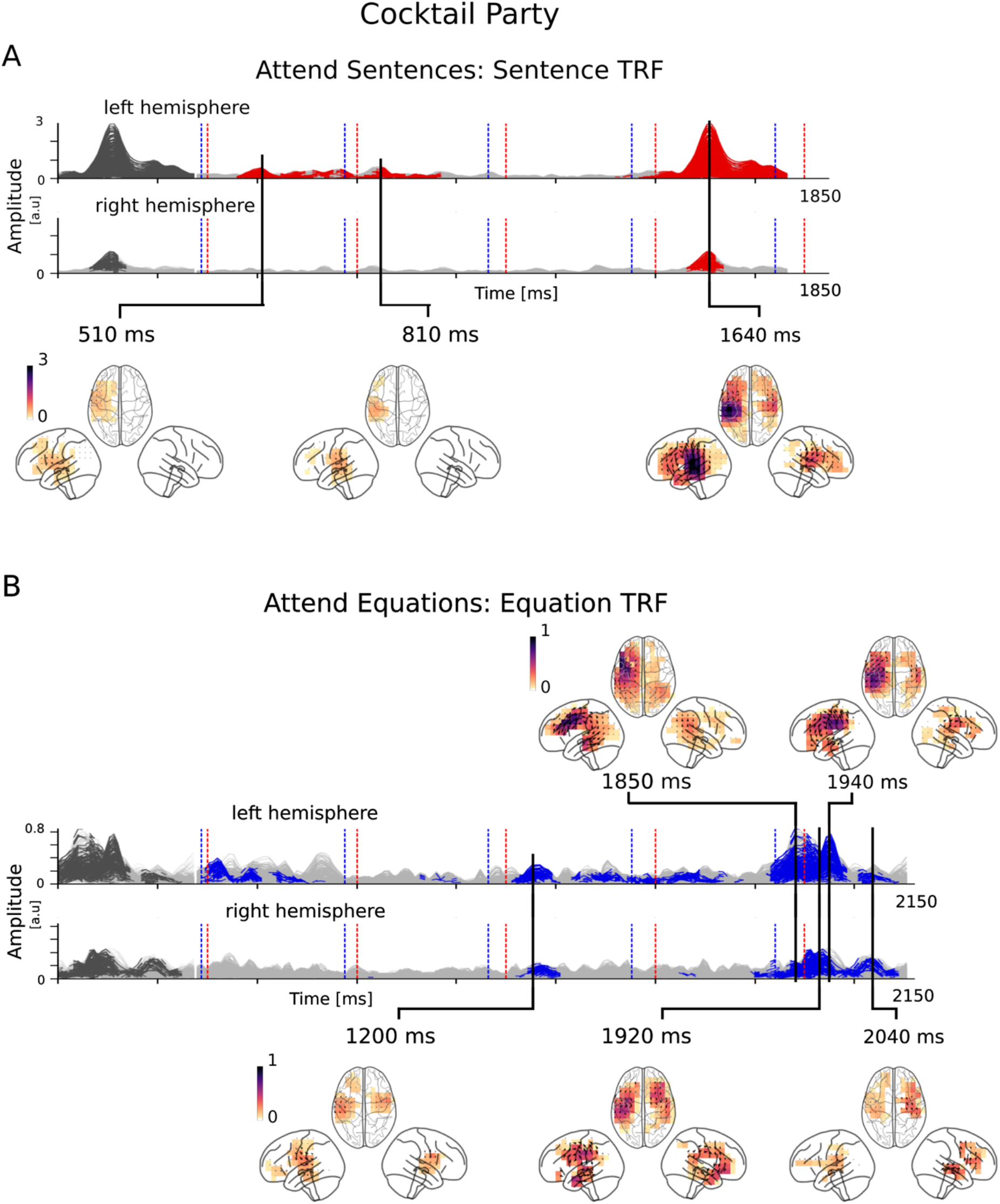
TRFs in the cocktail party conditions. Overlay plots of the TRF for each voxel averaged over subjects are shown as those in Fig. 5. Word and symbol onset times are shown in red and blue dashed lines respectively and are marked in both sentence and equation TRFs since both stimuli were present in the cocktail party conditions; again, it can be seen that responses contributions associated with them have been successfully regressed out. Differences between sentence and equation TRFs arise at later time points, with sentence TRFs being predominantly near left temporal areas, while equation TRFs are in bilateral temporal, motor, and parietal regions. The entire sentence and equation TRFs are shown in Video 2.

#### Video 2. TRFs in the cocktail party conditions

The sentence and equation TRFs are shown from 350 ms until 350 ms past the end of the sentence or equation. Traces at significant time points are shown in red (sentence processing) or blue (equation processing). The distribution of cortical activity at each time point is shown below each TRF, with color scales fixed over each entire TRF.

### 3.5. Decoder analysis

To further help differentiate between the cortical processing of equations and sentences, two types of linear decoders were trained on neural responses. 1) Classifiers at each time point that learned weights based on the MEG sensor topography at that time point. 2) Classifiers at each voxel that learned weights based on the temporal dynamics of the response at that voxel. The former was used to contrast the dynamics of equation and sentence processing (Fig. 7A). For the single speaker conditions, all time points showed significant decoding ability across subjects (*t_max_* = 11.3, *p_min_* < 0.001), with higher prediction success (as measured by *AUC*) at longer latencies. For the cocktail party conditions, decoding ability was significantly above chance only at longer latencies (*t_max_* = 6.45, *p_min_* < 0.001). While subjects listened to the equations, the identity of the arithmetic operator (e.g., ‘plus’ vs. ‘times’ or ‘less’) was reliably decoded from the MEG sensor topography during the time points when the operator and the subsequent operand were presented (Fig. 7B). Note that decoding accuracy was significantly above chance for time points ∼250-300 ms after the offset of the operator. This is considerably late for decoding based on mere auditory responses to acoustic features of the operator.

**Figure 7.**
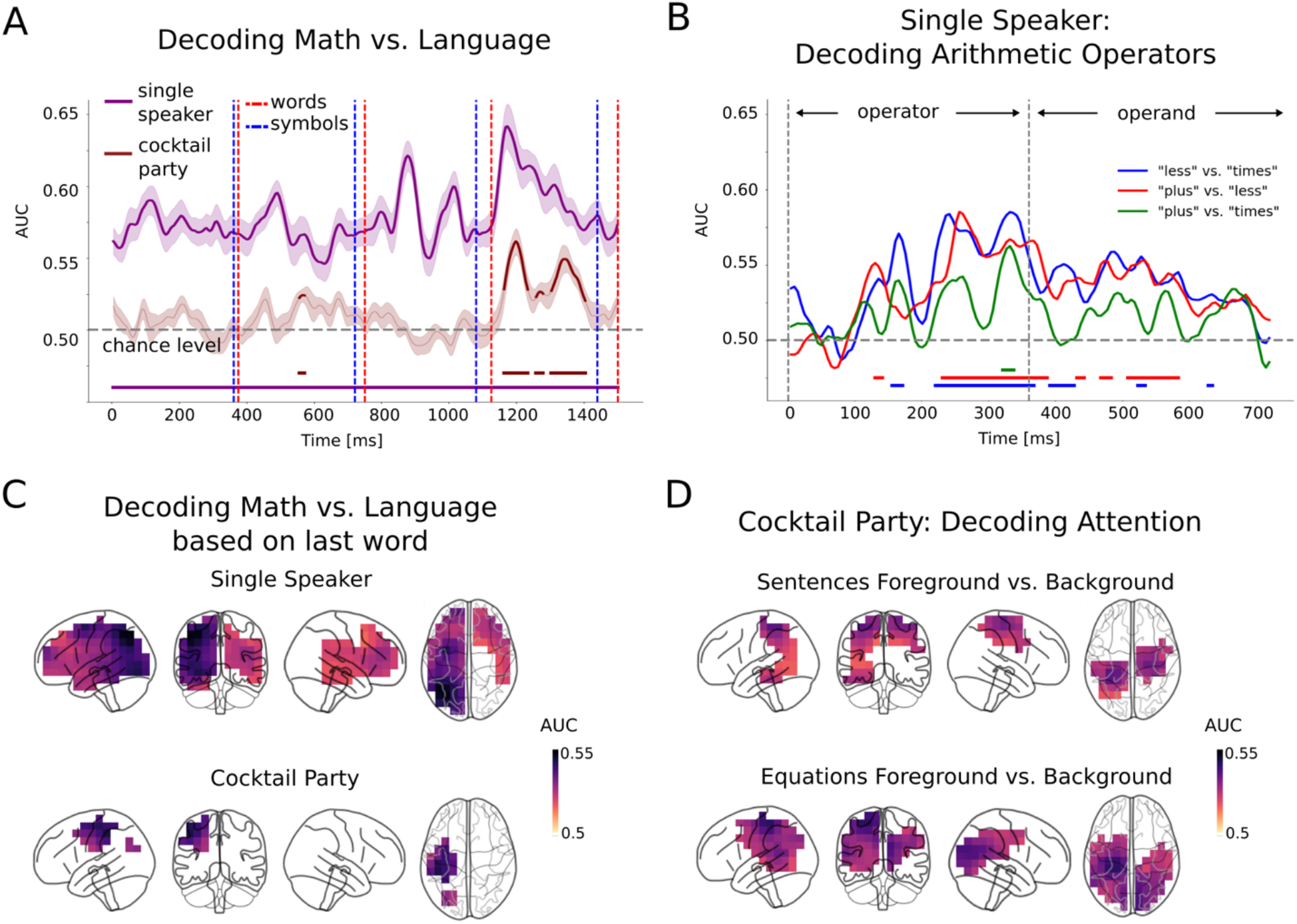
Decoding arithmetic and language processing. **A.** Performance of decoding attention condition (math vs. language) at each time point using MEG sensors for single speaker (purple) and cocktail party (brown). Prediction success is measured by *AUC*, which is plotted (mean and s.e.m. across subjects); time points where predictions are significantly above chance are marked by the horizontal bars at the bottom (every time point is significantly above chance for the single speaker case). The word and symbol onsets are also shown, and the decoding performance increases towards the end of the time window. **B.** Decoding arithmetic operators from sensor topographies. The time window of the operator and the subsequent operand was used for the 3 types of decoders. Time intervals where predictions are significantly above chance are marked by the colored horizontal bars at the bottom: all 3 operator comparisons could be significantly decoded. **C.** Decoding math vs. language based on the last word. During the single speaker conditions, most of the brain is significant. However, for the cocktail party conditions, more focal significant decoding is seen in IPS and superior parietal areas. Decoding based on the first word resulted in similar results (not shown). **D.** Decoding attention in the cocktail party conditions (*AUC* masked by significance across subjects). The sentence responses in foreground and background were decoded in left middle temporal and bilateral superior parietal areas. The equation responses in foreground and background were decoded in bilateral parietal areas.

Decoders at each voxel were also trained to differentiate attention to equations vs. sentences, based on the dynamics of the response at that voxel during the first or last words (Fig. 7C). The prediction success (as measured by *AUC*) was significant for large areas in the single speaker conditions both for first words (*t_max_* = 5.1, *p_min_* < 0.001) and last words (*t_max_* = 5.4, *p_min_* < 0.001) decoders. The *AUC* for first words decoders was significant for all regions in the left hemisphere except for areas in the inferior and middle temporal gyrus and all regions on the right hemisphere except the occipital lobe. The *AUC* for last words was significant for all regions in the left hemisphere and parts of frontal temporal lobes in the right hemisphere. For the cocktail party conditions the source-localized regions of significant prediction success were much more focal: the *AUC* was significant only in the IPS and superior parietal areas for both first words (*t_max_* = 5.3, *p_min_* < 0.001) and last words (*t_max_* = 4.3, *p_min_* = 0.014) decoders. These results suggest that the activity of voxels in left IPS and superior parietal areas most useful for discriminating between attending to equations vs. sentences. Finally, decoders at each voxel were also trained to decode the attention condition (foreground vs. background) from the response to the entire sentence or equation (Fig 7D). The *AUC* was significant in bilateral parietal areas for decoding whether arithmetic was in foreground vs. background (*t_max_* = 5.2, *p_min_* < 0.001), consistent with areas involved in arithmetic processing. For decoding whether language was in foreground vs. background, the *AUC* was significant (*t_max_* = 5.1, *p_min_* = 0.002) in left middle temporal areas (consistent with higher level language processing) and bilateral superior parietal areas (consistent with attention networks that are involved in discriminating auditory stimuli). Therefore, the decoding analysis is able to detect different cortical areas that may be involved in attention to language and arithmetic.

## 4 Discussion

Specific cortical patterns were found to be consistent across different analysis methods (frequency domain, TRFs, decoders) as illustrated in schematic Fig. 8. Sentence responses consistently localize to left temporal areas. Equation responses consistently show bilateral parietal activity, but also vary across analysis method (e.g., motor activity in TRFs). This may be due to different mechanisms involved in equation processing, although further investigation would be needed to support this claim.

**Figure 8:**
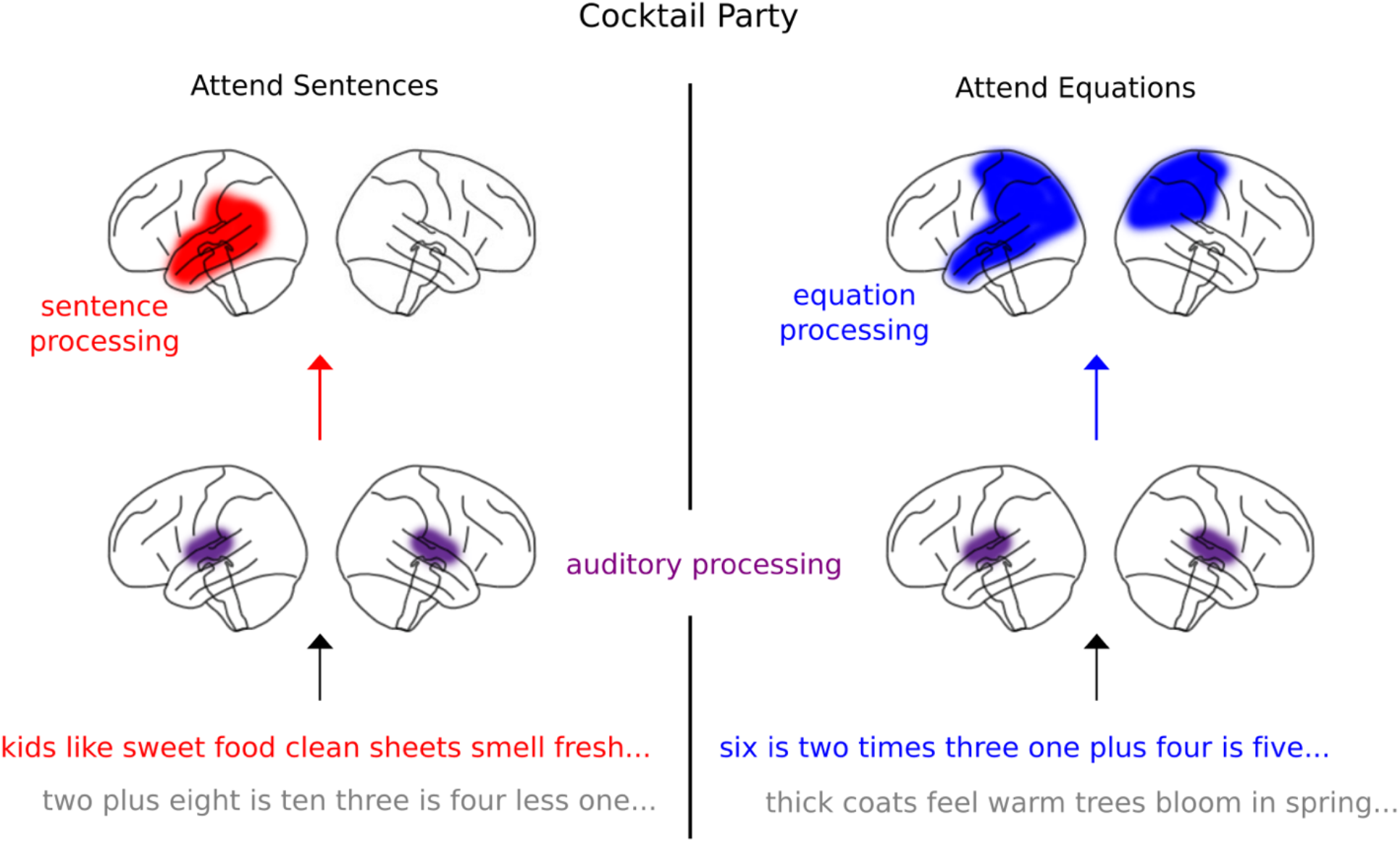
Schematic of cortical processing of sentences and equations. A schematic representation of sentence and equation processing is shown. Exemplars of both foreground and background of stimuli are shown at the bottom. The areas that were most consistent across all analysis methods (frequency domain, TRFs and decoders) are shown.

### 4.1. Sentence and equation rate responses

As expected, MEG responses to acoustic features source-localize to bilateral auditory cortex, and sentence rate responses source-localize to left temporal cortex, consistent with speech and language areas (Friederici, 2002, 2011; Vandenberghe et al., 2002; Hickok and Poeppel, 2007; Binder et al., 2009), similar to prior isochronous speech studies (Sheng et al., 2018). In contrast, we find equation rate responses localize to left parietal, temporal, and occipital areas. Arithmetic processing can activate IPS and parietal (Dehaene et al., 2003), angular gyrus (Göbel et al., 2001), temporal (Tang et al., 2006) and even occipital areas (Maruyama et al., 2012; Harvey and Dumoulin, 2017), perhaps due to internal visualization (Zago et al., 2001). Equation responses also localize to right temporal and parietal areas in cocktail party conditions, confirming that arithmetic processing is more bilateral than language processing (Dehaene and Cohen, 1997; Amalric and Dehaene, 2018, 2019). Critically, ANOVA analysis indicated that attended equation and sentence responses are significantly different. Unexpectedly, significant neural responses at the unattended sentence and equation rates were found in smaller temporal (consistent with language processing) and parietal (consistent with arithmetic processing) areas respectively. Some subjects may have been unable to sustain attention to the instructed stream for the entirety of this diotic stimulus and so briefly switched their attentional focus.

### 4.2. Left hemispheric dominance of equation responses

Equation responses were left dominant in both single speaker and cocktail party conditions. This could reflect left lateralized language processing since equations were presented using speech. However, arithmetic processing may also show left dominance (Pinel and Dehaene, 2009), perhaps due to precise calculations (Dehaene, 1999; Pica et al., 2004) or arithmetic fact retrieval (Dehaene et al., 2003; Grabner et al., 2009). These fast-paced stimuli required rapid calculations, and may have resulted in increased reliance on rote memory, which activates left hemispheric areas (Campbell and Austin, 2002). Specific strategies employed for calculation may also result in left lateralization—multiplication of small numbers is often performed using rote memory (Delazer et al., 1999; Ischebeck et al., 2006; Fehr et al., 2007), while subtraction is less commonly performed using memory and shows more bilateral activation (Schmithorst and Brown, 2004; Prado et al., 2011). Addition may recruit both these networks, depending on specific strategies utilized by individuals (Arsalidou and Taylor, 2011). We found no significant differences in equation responses when separated by operation type, perhaps because of individual variation in procedural calculation or retrieval strategies within the same operation (Tschentscher and Hauk, 2014). However, operation types were successfully decoded from the overall MEG signals (Fig. 8B), consistent with prior work (Pinheiro-Chagas et al., 2019), though this effect was not as robust as decoding stimulus type or attention. Overall, left hemispheric dominance of equation responses is supported by a combination of speech processing, precise calculations, and arithmetic fact retrieval.

### 4.3. Cortical correlates of behavioral performance

Neural responses to sentence, equation, word, and symbol rates were correlated with performance in detecting deviants, consistent with language-only isochronous studies (Ding et al., 2017). Sentence responses correlated with behavior in language areas, such as left auditory cortex, superior and middle temporal lobe and angular gyrus (Price, 2000; Binder et al., 2009; Karuza et al., 2013) in both single speaker and cocktail party conditions. In contrast, equation responses correlated with behavior in cocktail party conditions in posterior parietal areas, which are known to predict competence and performance in numerical tasks (Grabner et al., 2007; Lin et al., 2012, 2019; Lasne et al., 2019). The lack of significant behavioral correlations for equation responses in single speaker conditions may be due to several subjects performing at ceiling; equations had a restricted set of only 14 unique symbols, and the presence of the ‘is’ symbol in every equation might be structurally useful in tracking equation boundaries. Unexpectedly, behavioral correlations were also found for background symbol rate responses in parietal and occipital areas (and for background word rate responses in a small parietal region). Some studies show that acoustic features of background speech may also be tracked (Fiedler et al., 2019; Brodbeck et al., 2020). Since background word and symbol rates are present in the stimulus acoustics, increased effort or attention could enhance both behavioral performance and auditory responses at these rates. Representations of the background could enhance attentional selectivity in challenging cocktail party environments as suggested by Fiedler et al., 2019. However, note that sentence and equation responses were only correlated with behavior when attended. Overall, behavioral correlations in temporal and parietal regions suggest that these responses may reflect improved comprehension due to neural chunking of speech structures or successful calculations (Blanco-Elorrieta et al., 2019; Chen et al., 2020; Jin et al., 2020; Kaufeld et al., 2020; Teng et al., 2020).

### 4.4. Dynamics of arithmetic and language processing

TRFs for equation and sentence onsets were jointly estimated along with speech envelopes and word/equation onsets in order to regress out auditory responses, analogous to prior work with linguistic and auditory TRFs (Brodbeck et al., 2018a, 2018b; Broderick et al., 2018). Isochronous speech studies have found slow rhythmic activity (Zhang and Ding, 2017), which does not appear in our TRFs, perhaps due to implicit high-pass filtering (boosting favors sparse TRFs). Instead, we find large TRF peaks at sentence/equation boundaries. Prior studies have found late evoked responses specific to numbers and equations (Avancini et al., 2015). Large peaks appear in both sentence and equation TRFs 410–600 ms after the onset of the last word/symbol and may reflect processing of the completion of the sentence/equation. Sentence TRF peaks localize to left temporal areas, while equation TRF peaks show activity in bilateral parietal and temporal areas involved in numerical processing (Abd Hamid et al., 2011; Amalric and Dehaene, 2018), and motor areas, perhaps reflecting procedural calculation strategies (Tschentscher and Hauk, 2014). The latencies of these peaks are similar to prior arithmetic ERP studies (Iguchi and Hashimoto, 2000; Iijima and Nishitani, 2017). These sentence and equation TRF peaks may reflect several mechanisms; both shared (language processing, decision making), and separate (semantic vs. arithmetic processing), and further work is needed to disentangle these mechanisms. Finally, the cortical patterns of TRF peaks show more differences in the cocktail party than the single speaker conditions, suggesting that selective attention focuses the underlying cortical networks.

### 4.5 Decoding equation and sentence processing

Numbers and arithmetic operations have been previously decoded from cortical responses (Eger et al., 2009; Pinheiro-Chagas et al., 2019). In this study, the attended stimulus type (sentences or equations) was reliably decoded in single speaker conditions in broad cortical regions, perhaps due to highly correlated responses across cortex for this task. In contrast, decoding accuracy during cocktail party conditions was significant in left IPS and superior parietal areas suggesting that these regions are most important for discriminating between arithmetic and language processing. Both the attend-equations and the attend-sentences states could be decoded from bilateral superior parietal areas. This could be due to general attentional networks in fronto-parietal areas, or attentional segregation of foreground and background speech based on pitch or gender (Hill and Miller, 2010; Kristensen et al., 2013). Additionally, decoding the attend-equations state was also significant in bilateral parietal areas, consistent with arithmetic processing, while decoding the attend-sentences state was also significant in left middle temporal lobe, consistent with language processing (Hickok and Poeppel, 2007). Overall, MEG responses contain enough information to decode arithmetic vs. language processing, selective attention, and arithmetic operations.

### 4.6 The cocktail party paradigm highlights distinct cortical processes

Differences in source-localized sentence and equation responses were more prominent in the cocktail party than in the single speaker conditions for all analyses (frequency domain, TRFs and decoders). Responses to both stimuli presented simultaneously may have helped control for common auditory and pre-attentive responses. Abd Hamid et al., (2011) found fMRI activation in broader areas for spoken arithmetic with a noisy background than in quiet, perhaps due to increased effort. However, in our case, the background stimulus was not white noise, but rather meaningful non-mathematical speech. Our TRF analysis, which regresses out responses to background speech, as well as the selective attention task itself, may highlight specific cortical processes that best separate the arithmetic and language stimuli.

In summary, neural processing of spoken equations and sentences involves both overlapping and non-overlapping cortical networks. Behavioral correlations suggest that these neural responses may reflect improved comprehension and/or correct arithmetic calculations. Selective attention for equations focuses activity in temporal, parietal occipital and motor areas, and for sentences in temporal and superior parietal areas. This cocktail party paradigm is well suited to highlight the cortical networks underlying the processing of spoken arithmetic and language.

## Acknowledgements

This work was supported by DARPA (N660011824024), the National Science Foundation (SMA-1734892 and DGE-1449815), and the National Institutes of Health (R01-DC014085). The views, opinions and/or findings expressed are those of the authors and should not be interpreted as representing the official views or policies of the Department of Defense or the U.S. Government.

